# IDH1 and IDH2 mutants identified in cancer lose inhibition by isocitrate because of a change in their binding sites

**DOI:** 10.1101/425025

**Authors:** Juan P. Bascur, Melissa Alegría-Arcos, Ingrid Araya-Durán, Ezequiel I. Juritz, Fernando D. González-Nilo, Daniel E. Almonacid

## Abstract

IDH1 and IDH2 are human enzymes that convert isocitrate (ICT) into α-ketoglutarate (AKG). However, mutations in positions R132 of IDH1 and R140 and R172 of IDH2 cause these enzymes to convert AKG into 2-hydroxyglutarate (2HG). Concurrently, accumulation of 2HG in the cell is correlated with the development of cancer. This activity change is mainly due to the loss of the competitive inhibition by ICT of these enzymes, but the molecular mechanism behind this loss of inhibition is currently unknown. In this work we characterized the inhibition and loss of inhibition of IDH1 and IDH2 by means of the binding energies derived from molecular docking calculations. We characterized the substrate binding sites and how they differ among the mutant and wild type enzymes using a Jaccard similarity coefficient based on the residues involved in binding the substrates. We found that molecular docking effectively identifies the inhibition by ICT in the wild type and mutant enzymes that do not appear in tumors, and the loss of inhibition in the mutant enzymes that appear in tumors. Additionally, we found that the binding sites of the mutant enzymes are different among themselves. Finally, we found that the regulatory segment of IDH1 plays a prominent role in the change of binding sites between the mutant enzymes and the wild-type enzymes. Our findings show that the loss of inhibition is related to variations in the enzyme binding sites. Additionally, our findings show that a drug capable of targeting all IDH1 and IDH2 mutations in cancer is unlikely to be found due to significant differences among the binding sites of these paralogs. Moreover, the methodology developed here, which combines molecular docking calculations with binding site similarity estimation, can be useful for engineering enzymes, for instance, when aiming to modify the substrate affinity of an enzyme.

## Introduction

Human enzymes isocitrate dehydrogenase 1 (IDH1) and isocitrate dehydrogenase 2 (IDH2) are members of the NADP^+^ dependent isocitrate dehydrogenase family of enzymes. IDH1 is found in the cytoplasm, while IDH2 is found in the mitochondria. Both enzymes convert isocitrate (ICT) and NADP^+^ into α-ketoglutarate (AKG), CO2 and NADPH, refilling the cell NADPH reserves. IDH1 also catalyzes the reverse reaction during hypoxic conditions, refilling the ICT reserves. The active form of these enzymes is a homodimer presenting 2 active sites in the interface between the subunits. These enzymes present an open form when binding just NADP+ or NADPH, adopting the closed form upon binding ICT or AKG. It is well established that mutations in positions IDH1 R132, IDH2 R140 and IDH2 R172 are associated with cancer development, particularly in glioblastomas, lymphomas and leukemias (Yan et al., 2009; Marcucci et al., 2010; Cairns et al., 2012). These mutations confer the enzymes the activity of converting AKG and NADPH into 2-hydroxyglutarate (2HG) and NADP^+^, impeding their normal activity (Dang et al., 2009). Furthermore, accumulation of 2HG in the cell is strongly correlated with the development of cancer (Dang et al., 2009; Losman & Kaelin, 2013).

In a healthy cell the conversion of AKG into ICT is inhibited in presence of ICT, as ICT and AKG compete for the same active site. When the concentration of ICT decreases, as during hypoxia, conversion of AKG into ICT is uninhibited (Wise et al., 2011; Filipp et al., 2012). Mutant enzymes, however, are not inhibited in presence of ICT (Pietrak et al., 2011). It is believed that this lack of inhibition allows these enzymes to accept AKG as a substrate (Pietrak et al., 2011). The mechanism by which the mutant enzymes convert AKG into 2HG, instead of ICT is still being investigated. It has been hypothesized that it could result from the conformational changes of the residues surrounding the active site; or even changes in the kinetic mechanism of the mutant enzymes (Dang et al., 2009; Rendina et al., 2013). However, the mechanism by which the mutant enzymes lose their inhibition has not been investigated.

In this work, we present a detailed characterization of this inhibition loss mechanism through molecular docking simulations. We worked for that end with the structures of both mutant and wild type IDH1 and IDH2 enzymes and analyzed the binding energies and substrate binding sites. A novel methodology introduced in this study is the comparison of binding sites by computing a Jaccard similarity coefficient (Jaccard, 1912) based on the residues involved in substrate binding. We found that the binding energies were coherent with the observed loss of inhibition in the mutant enzymes. We also found that the binding sites present significant differences among the mutants and, in addition, evidence that some of these binding sites are functional.

## Materials and methods

### Sequence information

The nucleotide and amino acid sequences as well as SNP information for human IDH1 and IDH2, porcine IDH2 and *Escherichia coli’s* isocitrate dehydrogenase (IDH) were retrieved from the NCBI databases (NCBI Resource Coordinators, 2016) (Table 1). Mutations reported in tumours were obtained from the “Catalog of somatic mutations in cancer” (COSMIC) version 80 (Forbes et al., 2015). Only substitutions with both wild type and mutant nucleotides explicitly reported were included.

**Table 1.** Human IDH1, human IDH2, porcine IDH2 and Escherichia coli’s IDH reference sequences and SNPs. The only SNP registered for the IDH enzymes is located in position IDH1 R100.

| Gene | mRNA sequence | Protein sequence | SNP mRNA sequence | Reference SNP cluster |
| --- | --- | --- | --- | --- |
| Human IDH1 | NM_005896.3 | NP_005887.2 | NM_001282386.1 | rs777129475 |
| Human IDH2 | NM_002168.3 | NP_002159.2 | - | - |
| Porcine IDH2 | - | NP_001157479.1 | - | - |
| *E. coli* IDH | - | NP_415654.1 | - | - |

The catalytic residues of IDH1 and IDH2 were mapped from the *E. coli* IDH sequence by alignments performed by PROMALS3D (Pei, Kim & Grishin, 2008). The *E. coli* isocitrate dehydrogenase (IDH) catalytic residues were obtained from the entry M0007 of the Mechanism, Annotation and Classification in Enzymes (MACiE) database (Holliday et al., 2012). The regulatory segment, clefts flanking residues and the protein domains of human IDH2 and porcine IDH2 were mapped from human IDH1 by sequence alignment with PROMALS3D.

### Protein structures selection

We used the wild type enzymes as well as mutations reported in tumors for positions IDH1 R100, IDH1 R132, IDH2 R140 and IDH2 R172 in the COSMIC database to establish which mutant structures to use. The most common mutation for each site was selected and thus the structures of IDH1 R100Q, IDH1 R132H, IDH2 R140Q and IDH2 R172K were used.

The protein structures were either obtained from the Protein Data Bank (PDB) (Berman et al., 2000), from homology modeling using Modeller 9.17 (Šali & Blundell, 1993) combined with the Maestro Prime tool (Jacobson et al., 2004) or from modeled mutations using the Maestro Suite Platform (Schrödinger, 2015) and Maestro Prime tool (Table 2).

**Table 2.**
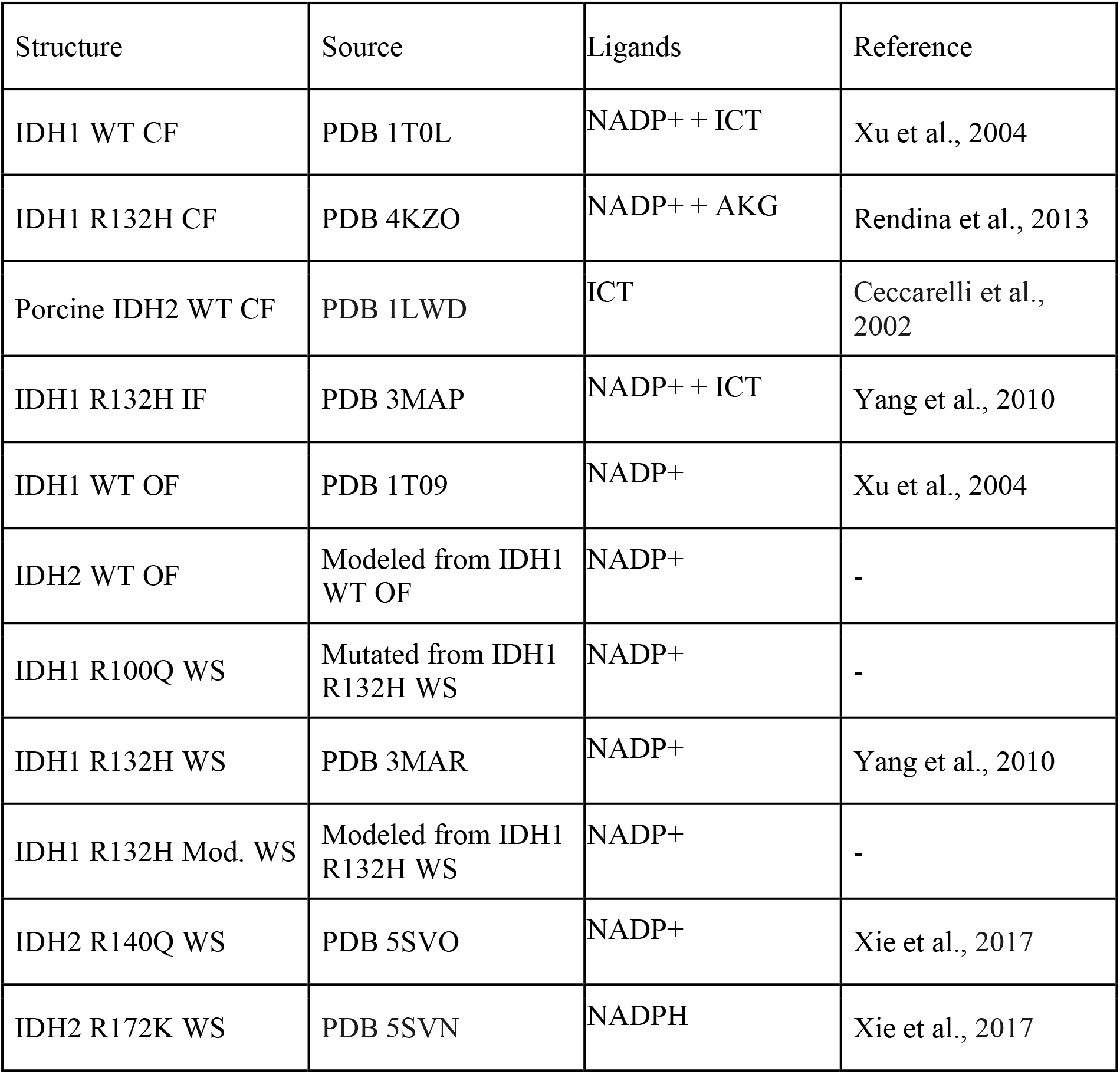
IDH1 and IDH2 WT and mutant enzymes studied in this work. The table includes the source of the structures (obtained from PDB, from homology modeling or from modeled mutations), ligands present in the structures and literature references.

| Structure | Source | Ligands | Reference |
| --- | --- | --- | --- |
| IDH1 WT CF | PDB 1T0L | NADP+ + ICT | Xu et al., 2004 |
| IDH1 R132H CF | PDB 4KZO | NADP+ + AKG | Rendina et al., 2013 |
| Porcine IDH2 WT CF | PDB 1LWD | ICT | Ceccarelli et al., 2002 |
| IDH1 R132H IF | PDB 3MAP | NADP+ + ICT | Yang et al., 2010 |
| IDH1 WT OF | PDB 1T09 | NADP+ | Xu et al., 2004 |
| IDH2 WT OF | Modeled from IDH1 WT OF | NADP+ | - |
| IDH1 R100Q WS | Mutated from IDH1 R132H WS | NADP+ | - |
| IDH1 R132H WS | PDB 3MAR | NADP+ | Yang et al., 2010 |
| IDH1 R132H Mod. WS | Modeled from IDH1 R132H WS | NADP+ | - |
| IDH2 R140Q WS | PDB 5SVO | NADP+ | Xie et al., 2017 |
| IDH2 R172K WS | PDB 5SVN | NADPH | Xie et al., 2017 |

IDH1 WT form is open when bound to NADP+ (Xu et al., 2004) and closed when it binds NADP+ and ICT (Xu et al., 2004). On the other hand, the form of IDH1 R132H is open when binding NADP+ (Yang et al., 2010), intermediate when it binds NADP+ and ICT (Yang et al., 2010) and closed when is bound to NADPH and AKG (Rendina et al., 2013). A similar behavior is reported for IDH2 (Lv et al., 2012). The conformation from the forms of all studied structures were obtained from their original publications. Once the form was identified, it was annotated as either open form (OF), intermediate form (IF) or closed form (CF). Structures without identified form in their original publications also did not have ICT or AKG bound, so they were annotated as without substrate (WS) to highlight this common feature. IDH1 WT OF (PDB ID: 1T09) and the modeled structure IDH2 WT OF did not have substrates bound, but they were annotated as OF because the form of IDH1 WT OF is explicitly stated in its original publication. The form of the modeled structures was annotated according to their template.

#### Closed form structures

Human IDH1 WT CF structure was downloaded from the PDB database (PDB ID: 1T0L). Porcine IDH2 CF (PDB ID: 1LWD) was used as a substitute for human IDH2 WT CF, as there are no structures for the latter deposited in the PDB database. IDH2 WT CF was not modeled as it was not required for the molecular docking simulations. The only closed form of the mutant enzymes found in the PDB was IDH1 R132H CF (PDB ID: 4KZO), and it was defined as the representative structure for all closed forms of the mutant enzymes.

#### Intermediate form structures

The only intermediate form available in the PDB database was IDH1 R132H IF (PDB ID: 3MAP) and, therefore, it was defined as the representative structure for all intermediate forms.

#### Open form structures

IDH1 WT OF was available in the PDB database (PDB ID: 1T09). IDH2 WT OF was not in the PDB, so it was modeled based on the sequence of IDH2 WT and the structure of IDH1 WT OF due to the high amino acid identity between IDH1 and IDH2 (>70%). The model was generated using Modeller, and the model quality was estimated using the DOPE score of Modeller. The NADP+ cofactor was added to the model by structural superposition with the template. The protein-cofactor complex structure was minimized using the Prime tool of Maestro in order to keep the reported interactions. The mutant open form structures were not modeled, as they were expected to be among the set of structures without substrate.

#### Structures without substrate

In the IDH1 R132H WS structure obtained from the PDB (PDB ID: 3MAR) some of the unresolved residues were considered relevant for the protein-ligand molecular docking due to their possible proximity to the binding site. An additional structure of IDH1 R132H WS with all its residues was modeled over the structure of IDH1 R132H WS and defined as IDH1 R132H Modeled (Mod.) WS. The protocol followed for modeling was the same as for IDH2 WT OF model. Because IDH1 R100Q WS was not available in PDB database, it was modeled by introducing mutations into the structure of IDH1 R132H WS. The structure was mutated (R100Q and H132R) with the Maestro Suite Platform on all chains and the mutated residues were minimized using the Prime tool of Maestro. IDH2 R140Q WS (PDB ID: 5SVO) and IDH2 R172K WS (PDB ID: 5SVN) were available in the PDB database.

We aimed to perform a comparison between the binding sites of the open, intermediate and closed structures among the WT and mutant IDH1 and IDH2 enzymes. The secondary structure of the regulatory segments was annotated using the Maestro Suite Platform (Table 3).

**Table 3.**
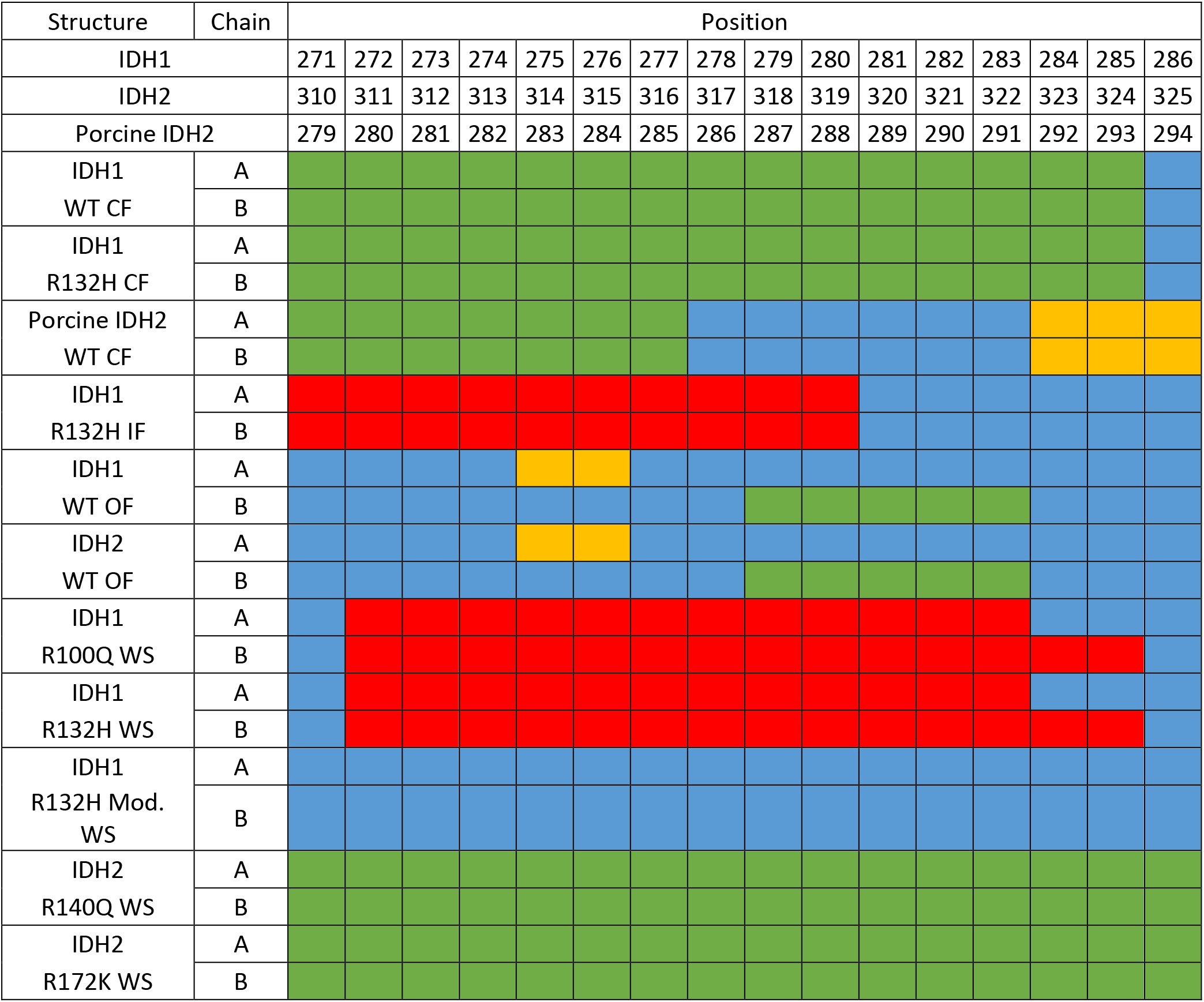
Secondary structure of the regulatory segment of IDH1 and IDH2 WT and mutant enzymes. The table shows the secondary structure for each residue of the regulatory segment. Green: α-helix, yellow: β-sheet, blue: loop, and red: unresolved residue.

### Structural characterization of the complexes

In the IDH1 structure the active site is located in a cleft (active site cleft) and behind another well-defined cleft (back cleft). In the open form the active site cleft is wider than the back cleft, while in the closed form the back cleft is wider. Moreover, in the open form the active site cleft of one subunit is wider than the cleft of the other subunit, and are accordingly defined as open cleft (OC) and semi-open cleft (SC) (Figure 1) (Xu et al., 2004). The open and semi-open clefts of the open form structures were identified by measuring the width of the active site cleft and the back cleft of each subunit. This step was essential to identify the OC and the SC of each structure, so the binding site and binding energies for the complexes were compared among clefts of the same kind. The residues flanking the clefts in IDH1 were obtained from the literature (Xu et al., 2004) and the residues in IDH2 and porcine IDH2 were inferred by sequence alignment with IDH1 (Table 4). The back cleft is flanked by residues of the same subunit, whereas the active site cleft is flanked by residues on opposite subunits, so we emphasized whether the flanking residues belonged to the same chain (S. Ch.) or the opposite chain (O. Ch.). The width was measured as the distance between the α-carbons of the flanking residues.

**Figure 1.**
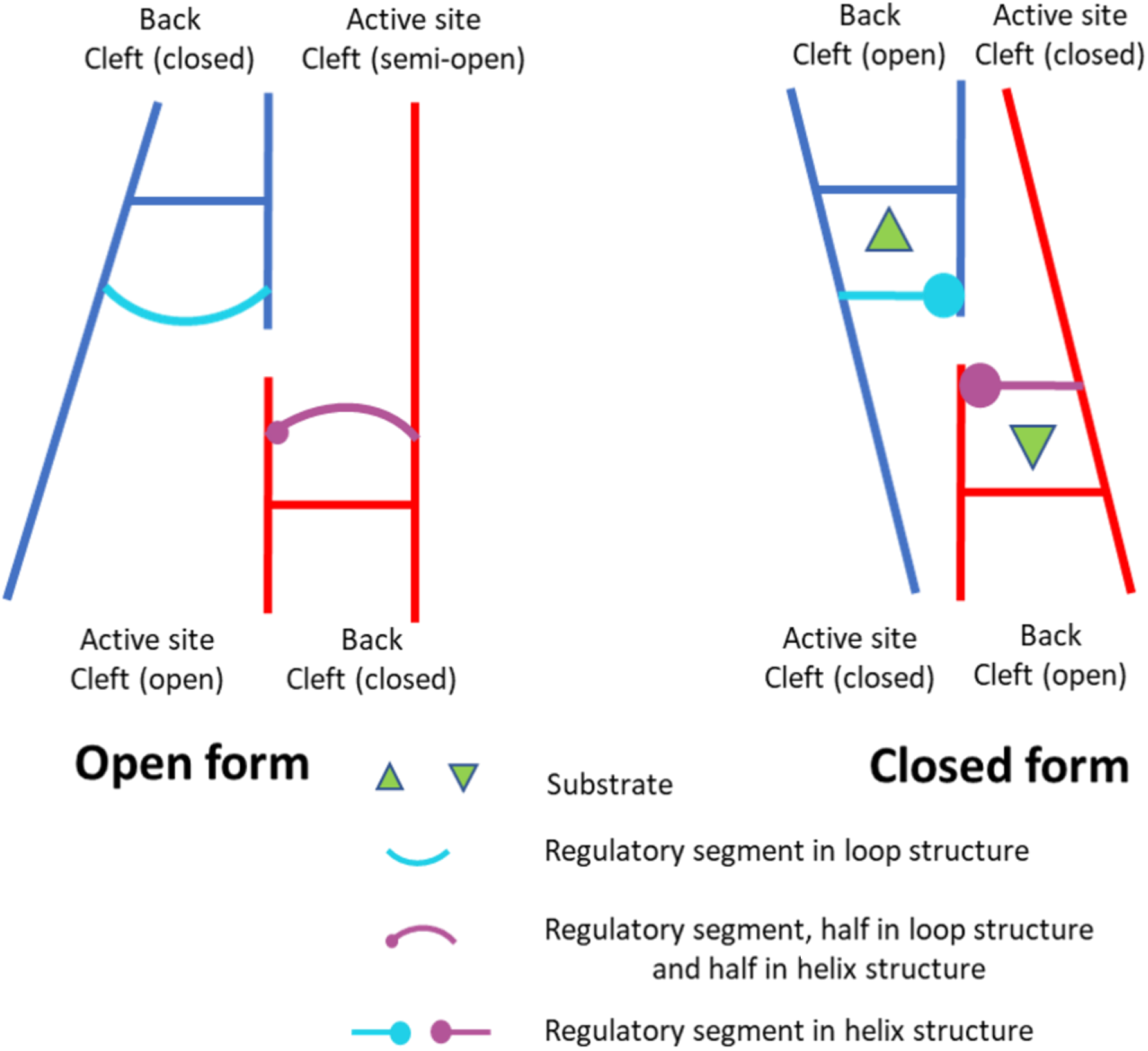
Relation between the width of the active site and back clefts and the form of the IDH enzymes. The blue and red lines represent the subunits of the dimer. The triangle represents the substrate. In the open form, the active site cleft is open and the back cleft is closed, and vice versa in the closed form. The width of the clefts is correlated to the secondary structure adopted by the regulatory segment. This figure is based on Figure 6 of the article of Xu et al., 2004.

**Table 4.** Residues flanking the active site cleft and the back cleft of IDH1 and IDH2. Note that the back cleft is flanked by residues of the same protein chain (S. Ch.), whereas the active site cleft is flanked by a residue of the S. Ch. and a residue of the opposite protein chain (O. Ch.).

| Enzyme | Active site cleft residues |  | Back cleft residues |  |
| --- | --- | --- | --- | --- |
| Human IDH1 | 76 S. Ch. | 250 O. Ch. | 199 S. Ch. | 342 S. Ch. |
| Human IDH2 | 116 S. Ch. | 289 O. Ch. | 238 S. Ch. | 381 S. Ch. |
| Porcine IDH2 | 85 S. Ch. | 258 O. Ch. | 207 S. Ch. | 350 S. Ch. |

Root Mean Square Deviation (RMSD) between both subunits of each structure was also calculated in order to classify the structures as symmetric or asymmetric. These measurements were used for later analyses of the binding sites. RMSD values were calculated based on the α-carbons of the structures using Protein3Dfit (Lessel & Schomburg, 1994).

### Molecular docking

A validation was necessary to prove the fitness of the molecular docking algorithms used for the enzyme-substrate complex system under study. The method for validating consisted of removing cocrystallized molecules from protein complex structures and then re-docking them on their original site, testing if the docked molecule position was equivalent to the original molecule position (Warren et al., 2006). The molecular docking algorithms Autodock Vina (Morris et al., 2009) and Glide XP (Schrödinger, 2015) were validated by redocking ICT on IDH1 WT CF and AKG on IDH1 R132H CF (data not shown). Glide XP outperformed Autodock Vina and therefore was selected for the molecular docking simulations reported in this work.

Receptor files for the molecular docking simulations were prepared with the “Protein Preparation Wizard” of the Maestro Suite Platform. All substrate structures were obtained from PubChem (Kim et al., 2016). PubChem CID code for ICT is 1198 and for AKG is 51. Ligand files were prepared using the “Ligand Preparation Wizard” of the Maestro Suite Platform. The size of the grid was established at 40Åx40Åx40Å and it was centered on the centromere of the binding residues in the closed form of the enzymes, calculated independently for each grid (Table 5). For all structures, binding residues were defined as the residues within 4Å from the substrates, and the centromere was calculated using the α-carbons of the binding residues. The closed structure forms used to obtain the binding residues in closed formation were IDH1 WT CF and IDH1 R132H CF. The binding residues were identified in all chains of both structures and included in the residue set used for defining the centromere location. Interestingly, both closed structures had the same binding residues. The binding residues of IDH2 were mapped from IDH1 by sequence alignment.

**Table 5.** Human IDH1 and IDH2 closed form binding residues. The binding residues are the residues within 4Å from the substrates. IDH1 binding residues were obtained from IDH1 WT CF and IDH2 residues were mapped by sequence alignment with IDH1.

| Enzyme | Same chain |  |  |  |  |  |  |  |  | Opposite chain |  |  |  |
| --- | --- | --- | --- | --- | --- | --- | --- | --- | --- | --- | --- | --- | --- |
| Human IDH1 | 77 | 94 | 96 | 100 | 109 | 132 | 139 | 275 | 308 | 212 | 214 | 215 | 252 |
| Human IDH2 | 117 | 134 | 136 | 140 | 149 | 172 | 179 | 314 | 347 | 251 | 253 | 254 | 291 |

### Molecular docking comparisons

We used the docking score of Glide XP as the binding energies of the docking simulations. The differences registered among the binding energies of both ICT and AKG and the receptors in the molecular dockings were obtained in order to confirm that the mutant enzymes were, in effect, uninhibited when compared to the wild type enzymes (hereafter ΔΔG). The mutant ΔΔG value was then compared with the registered difference in the wild type enzymes (hereafter ΔΔΔG). The lowest registered ΔΔG value of the WT enzymes for each cleft (LΔΔG) was used in the comparisons to increase the stringency of our analyses.

The binding sites of the molecular docking assays were defined as the residues within a 4Å distance from the substrates. The similarity between the binding sites was measured using the Jaccard Similarity Coefficient (JS) (Jaccard, 1912) as shown in Eq. 1. The similarity between the binding sites of the substrates was computed considering the similarity between the binding sites of both substrates (Both S.) ICT and AKG together, or each substrate individually. The structure of IDH1 R132H CF was ignored as it presents the same binding residues as IDH1 WT CF.

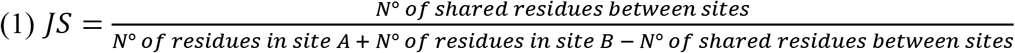

## Results

### Reported mutations for IDH1 and IDH2

Substitutions reported in tumours for positions IDH1 R100, IDH1 R132, IDH2 R140 and IDH2 R172 were considered for identifying the most common mutation per position (Table 6). A total of 10, 783, 5604 and 412 substitutions were found for each position respectively. The most common mutations were IDH1 R100Q, IDH1 R132H, IDH2 R140Q and IDH2 R172K. No proline substitutions were reported. It has been suggested that proline substitutions are unrelated to cancer due to their significant disruptive effect on protein structures (Pietrak et al., 2011).

**Table 6.** Substitutions of human IDH1 and IDH2 in cancer. Columns represent triplet nucleotides (in order) and rows registered substitutions on those nucleotides. The amino acidic product and the abundance of each nucleotide substitution is informed. IDH1 R100 substitutions are informed for the two known alleles.

| Subs. | IDH1 R100 Triplet |  |  | IDH2 R140 Triplet |  |  | IDH1 R132 Triplet |  |  | IDH2 R172 Triplet |  |  |
| --- | --- | --- | --- | --- | --- | --- | --- | --- | --- | --- | --- | --- |
|  | C, A | G | A | C | G | G | C | G | T | A | G | G |
| A | WT | Gln 8,<br>Lys 0 | WT | Arg<br>0 | Gln<br>753 | Arg<br>0 | Ser<br>165 | His<br>4371 | Arg<br>0 | WT | Lys<br>300 | Arg<br>1 |
| G | Gly<br>0 | WT | Arg 0,<br>Arg 0 | Gly<br>4 | WT | WT | Gly<br>196 | WT | Arg<br>0 | Gly<br>13 | WT | WT |
| T | STOP<br>2 | Leu 0,<br>Ile 0 | Arg 0,<br>Ser 0 | Trp<br>14 | Leu<br>12 | Arg<br>0 | Cys<br>769 | Leu<br>103 | WT | Trp<br>20 | Met<br>31 | Ser<br>29 |
| C | WT | Pro 0,<br>Thr 0 | Arg 0,<br>Ser 0 | WT | Pro<br>0 | Arg<br>0 | WT | Pro<br>0 | Arg<br>0 | Arg<br>1 | Thr<br>3 | Ser<br>14 |

### Clefts widths and symmetry of the structures

IDH1 and IDH2 are dimers with 2 active sites that are asymmetric in open form (inactive) and symmetric in closed form (active). The active site is inside a cleft (active site cleft) and behind a second cleft (back cleft). The asymmetry of the active sites is mainly due to the fact that one active site cleft is wider than the other (the open cleft and the semi-open cleft). In the open form, the active site opens and the back cleft closes, and the opposite happens when the protein adopts the closed form (Figure 1) (Xu et al., 2004).

We measured the cleft widths to determine if the enzyme form is closed or open (Table 7). We also identified in each structure the corresponding cleft (open or semi-open) for each active site cleft (the wider cleft was the open cleft). We used the values of IDH1 WT OF and IDH1 WT CF as references to determine if a structure is in open or closed form. Additionally, RMSD value between α-carbons of chains was measured to determine if the enzyme was symmetric or asymmetric.

**Table 7.** Cleft widths and RMSDs between chains of IDH1 and IDH2 WT and mutant enzymes [Å]. Blue cells indicate the width of the entrance of the active site cleft and the back cleft for each chain of the dimeric structures. Yellow cells indicate the difference among between the widths of the same clefts in opposite chains. Green cells report the RMSD values between both chains of the dimer structure. Darker colors highlight values of larger magnitude.

| Structure | Chain A | | Chain B | | $\Delta$ | | RMSD |
| --- | --- | --- | --- | --- | --- | --- | --- |
|  | Active site cleft | Back cleft | Active site cleft | Back cleft | Active site cleft | Back cleft |  |
| IDH1 WT CF | 12.8 | 16.2 | 12.6 | 16.2 | -0.2 | 0.0 | 0.3 |
| IDH1 R132H CF | 12.4 | 16.0 | 12.5 | 15.8 | 0.1 | -0.2 | 0.3 |
| Porcine IDH2 CF | 13.7 | 15.3 | 13.7 | 15.4 | 0.0 | 0.1 | 0.2 |
| IDH1 R132H IF | 19.6 | 10.7 | 17.5 | 11.0 | -2.1 | 0.3 | 0.6 |
| IDH1 WT OF | 21.2 | 10.9 | 18.8 | 11.7 | -2.4 | 0.8 | 0.7 |
| IDH2 WT OF | 21.4 | 10.8 | 19.2 | 11.7 | -2.2 | 0.9 | 0.8 |
| IDH1 R100Q WS | 20.5 | 11.6 | 18.3 | 11.3 | -2.2 | -0.3 | 0.6 |
| IDH1 R132H WS | 20.7 | 11.6 | 18.6 | 11.3 | -2.1 | -0.3 | 0.6 |
| IDH1 R132H Mod. WS | 20.7 | 11.6 | 18.6 | 11.3 | -2.1 | -0.3 | 0.6 |
| IDH2 R140Q WS | 21.6 | 12.3 | 21.5 | 12.1 | -0.1 | -0.2 | 0.4 |
| IDH2 R172K WS | 12.5 | 16.1 | 12.4 | 16.0 | -0.1 | -0.1 | 0.2 |

The reference values obtained for IDH1 WT OF and IDH1 WT CF were:

Open form: active site cleft widths 21 Å and 19 Å; back cleft widths 11 Å and 12 Å; difference between the active site clefts was −2 Å; difference between the back clefts was 1 Å; RMSD 0.7 Å.

Closed form: active site cleft widths 13 Å; back cleft widths 16 Å; no difference registered between clefts of the same kind; RMSD 0.3 Å.

By comparing the rest of the values in Table 7 with these reference values, we obtained the following results:

- All structures that were either OF or CF present values corresponding to their form.
- IDH1 R132H IF have values corresponding to an open form.
- All IDH1 WS mutant structures have distances and RMSD values that are closer to the open form than to the closed form.
- IDH2 R140Q WS: Its active site cleft and back cleft have widths that are similar in magnitude to those of an open form, but the differences between cleft widths of the same kind and RMSD between chains are similar to those of the closed form.
- IDH2 R172K WS: All values correspond to the closed form.

### Secondary structure of the regulatory segment

IDH1, and by homology IDH2, present a self-regulating mechanism involved in blocking the conversion of ICT into AKG when the concentration of ICT is low. The segment of the enzyme participating in this mechanism, the regulatory segment, blocks the access of the substrate to the catalytic residues, and can only be displaced when enough concentration of the substrate is reached (Xu et al., 2004; Zhao et al., 2009). In the open form, the regulatory segment forms a loop in the open cleft, while in the semi-open cleft the first half forms a loop but the second half adopts an α-helix structure. In the closed form, the totality of the regulatory segment adopts an α-helix structure in both active site clefts. Thus, secondary structure variations in the regulatory segment assist the enzyme changes from open into closed form and *vice versa* (Figure 1) (Xu et al., 2004).

Mutation IDH1 R132H increases the flexibility of the regulatory segment, hindering its regulatory function. This increased flexibility also prevents the regulatory segment from being resolved during X-ray crystallography, as it happens in the structure IDH1 R132H WS (Yang et al., 2010). As Dang et al exposed in 2008, the absence of the regulatory segment in the binding site forms a new binding site (Dang et al., 2009). Due to the importance of the secondary structure of the regulatory segment, we annotated the secondary structure of all its residues in all the studied structures (Table 3).

The following findings are worth considering:

- IDH1 R132H IF regulatory segment has some unresolved residues.
- IDH2 R140Q WS and IDH2 R172K WS regulatory segments present an α-helix structure, in concordance with the observations above that the differences between cleft widths of the same kind and RMSD values between chains are equivalent to those of the known closed forms.
- IDH1 R132H Mod. WS regulatory segment has a loop structure, as expected, given that it was modelled using the IDH1 R132H WS structure that had most of its regulatory segment structurally unresolved.

### Molecular docking binding energies

We docked both substrates ICT and AKG into the active sites of the open forms of the WT enzyme structures as well as in the WS forms of the mutant enzymes. Then we evaluated their binding energies (i.e. ΔG) in order to confirm that the binding energy differences between the substrates (ΔΔG) was smaller in the mutant enzymes (IDH1 R132H WS, IDH2 R140Q WS and IDH2 R172K WS) than in the wild type and IDH1 R100Q WS structures. Thus, explaining the loss of inhibition by ICT in the former group of mutants. We used the lowest binding energy of the wild type complexes (LΔΔG) for each cleft as reference, independent of the enzyme studied (IDH1 or IDH2), to increase the stringency of our calculations (Table 8). The lowest ΔΔG value, LΔΔG, was 1.8 Kcal/mol for the open cleft (from IDH2) and 1.0 Kcal/mol for the semi-open cleft (from IDH1). As expected, we found that the mutants that appear in tumors had a smaller ΔΔG than the wild types in at least one binding site. The sites (i.e. clefts) with smaller ΔΔG were IDH1 R132H WS OC, IDH2 R140Q WS SC and IDH2 R172K WS OC.

### Binding residues

We identified the residues in the binding sites that bind each substrate (Figure 2) and compared them using the Jaccard similarity index (Figure 3) to define a binding site similarity (BSS). We did not attempt to identify a BSS threshold above which sites are similar with statistical significance. Instead, we defined a threshold of 0.5 to differentiate two broad populations of comparisons, and just consider those with BSS greater than 0.5 in our discussion of results, noting that 15.6% of all comparisons made were in this group. We also aimed to identify binding sites among the mutant enzymes that were similar to the binding sites of known functional structures. These known functional structures were IDH1 WT CF, IDH1 R132H IF, IDH1 WT OF and IDH2 WT OF.

**Figure 2.**
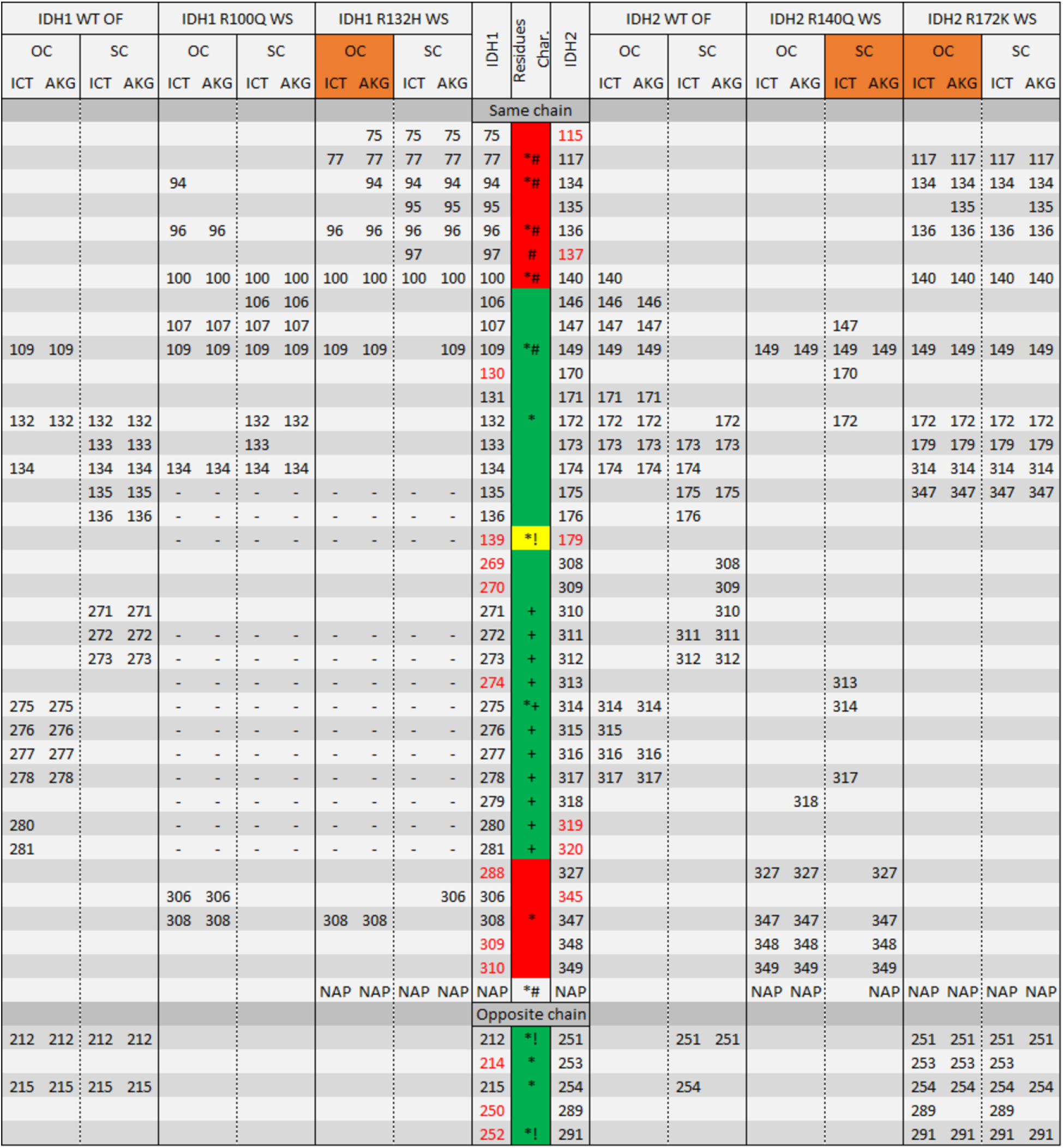
Residues binding ICT and AKG in the binding sites of IDH1 and IDH2 WT and mutant enzymes. The first row of the table indicates the structure, the second the cleft and the ligands. Orange cells indicate the mutant binding sites (i.e. clefts) with smaller ΔΔG compared to the WT ΔΔG, as reported in Table 8. The position of all the residues involved in substrate binding are indicated in the columns. Unresolved residues are indicated by an “-”. The three central columns show the position equivalences between IDH1 and IDH2, and serve as reference for the rest of the table. IDH1 residues are on the left columns, and IDH2 residues are on the right columns. Numbers in red in the IDH1 and IDH2 central columns indicate residues not involved in substrate binding in IDH1 or IDH2, respectively. The central column indicates structural and functional characteristics of the residues. The color of the central column indicates to which domain the residues belong. Red: Rossmann domain, green: α/β-sandwich domain, yellow: clasp domain. Symbols in the middle column stand for: “!” catalytic residue, “+” regulatory segment, “*” binding residue in the closed form, and “#” binding residue in the intermediate form.

We found that the BSS in IDH1 WT OF and IDH2 WT OF was higher for the semi-open cleft (BSS = 0.83) that for the open cleft (BSS = 0.44). We also found that all the mutant binding sites had a BSS lower than 0.5 with other mutant sites and were therefore considered each as unique. The following mutant binding sites presented significant similarities with one of the known functional binding sites:

- On both clefts, the binding sites in IDH1 R132H WS for both substrates (Both S.), as well as ICT and AKG separately, are similar to the binding sites in IDH1 R132H IF.
- On both clefts, the binding sites in IDH2 R172K WS for Both S., as well as ICT and AKG separately, are similar to the binding sites in IDH1 WT CF.

## Discussion

### Contrasts between IDH1 R100Q WS and IDH1 R132H WS binding sites

IDH1 R100Q WS structure was modelled by introducing two mutations (R100Q and H132R) on both chains of IDH1 R132H WS. Although small structural changes were expected for IDH1 R100Q WS and IDH1 R132H WS binding sites, significant variations were detected in both the binding energies and the binding residues among these structures. The binding energies obtained by our molecular dockings indicated that IDH1 R100Q conserves its inhibition by ICT, and thus it is not expected to produce 2HG. Although to our knowledge there are no functional characterizations of IDH1 R100Q, there is an IDH1 R100 mutant, IDH1 R100A, that has been proven to produce 2HG (Ward et al., 2012). However, nucleotide substitutions resulting in alanine mutations are rarely found in tumours and, in addition, it is not among the possible amino acid substitutions for IDH1 R100 position attained with just one nucleotide substitution. Instead, the most common mutation of IDH1 R100 in cancer is IDH1 R100Q (Table 6). This is due to the presence of a CpG site on IDH1 R100. It is well established that CpG (CG) sites are prone to mutating into TG sites (Cheng & Blumenthal, 2011). It is interesting to note that the two reported mutations in IDH1 R100 are CG -> TG in the sense (R100X) and antisense (R100Q) strands.

### The binding sites of the mutant enzymes are different among themselves

One of the expected results of this work was finding a high similarity between the binding sites of IDH1 R100Q WS and IDH2 R140Q WS and between those of IDH1 R132H WS and IDH2 R172K WS, given that mutations occur in analogous positions. However, our results show large differences among these binding sites, as showed in Figure 3. But even though binding sites are very different, all of these mutants, with the exception of IDH1 R100Q, produce 2HG. Even more, it has been reported that mutating IDH1 R132H analogous positions in isocitrate dehydrogenase enzymes leads to 2HG production in at least two yeast species (Song et al., 2014).

**Figure 3.**
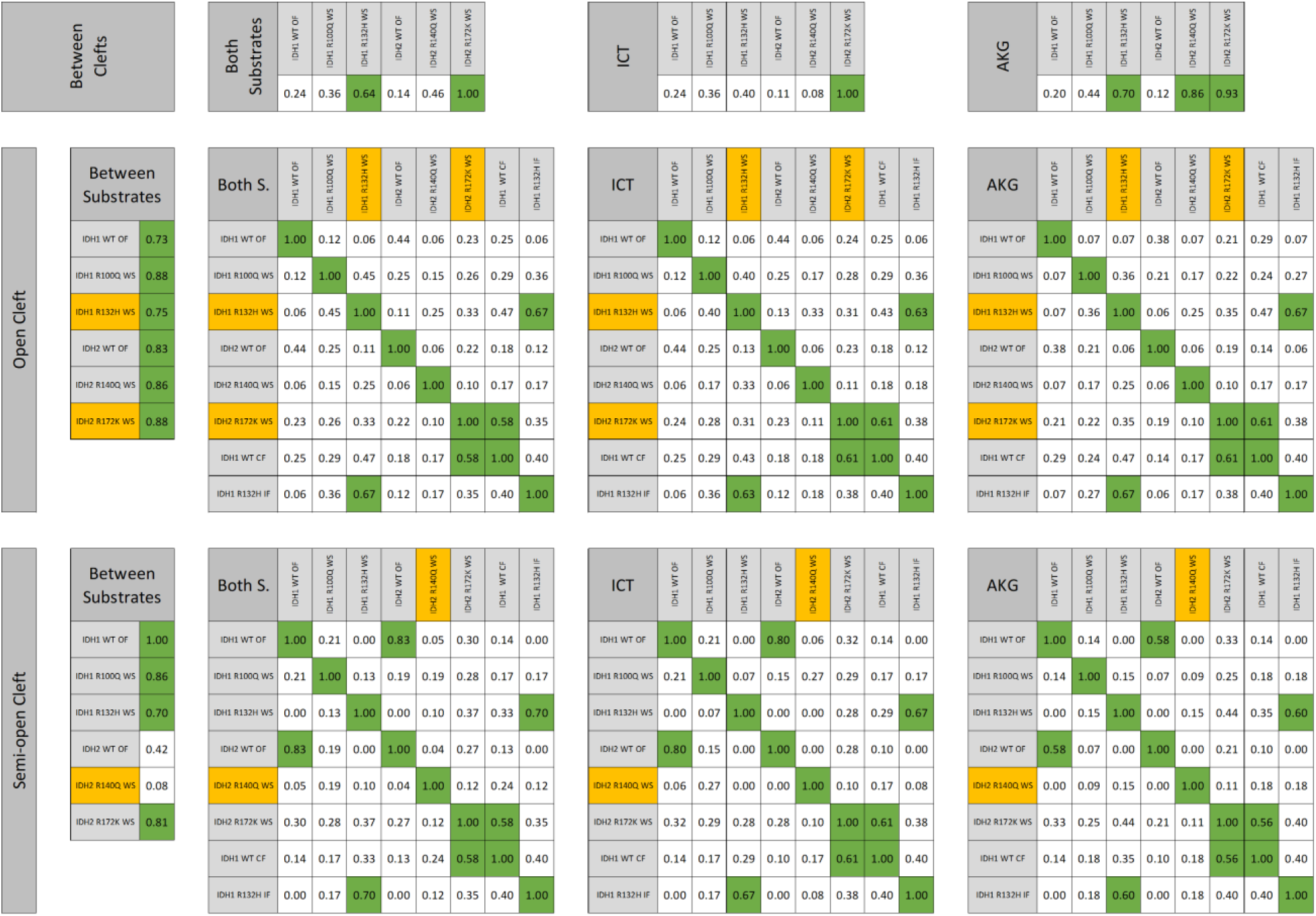
Similarities among binding sites of IDH1 and IDH2 WT and mutant enzymes. The similarity values were calculated using the Jaccard coefficient based on the residues involved in substrate binding. Orange cells indicate binding sites (i.e. clefts) with smaller ΔΔG compared to the WT ΔΔG, as reported in the Table 8. Green cells indicate Jaccard coefficient values over 0.50, an arbitrary threshold to highlight high similarity among sites. From top to bottom, the rows of tables describe similarities of the open cleft and the semi-open cleft in the same structure, only the open cleft and only the semi-open cleft. From left to right, the first tables describe the similarities between the binding site of both substrates in the same structure. The second tables describe the similarities between the binding sites of two structures when considering both substrates. The third and fourth tables are equivalent to the second table but considering only ICT or AKG, respectively.

Differences among mutants are registered not only in the binding sites but also in other structural features. The regulatory segment is unresolved in IDH1 R132H WS, so presumably it is behaving as a flexible loop. However, the regulatory segment in IDH2 R140Q WS and IDH2 R172K WS are fully folded into α-helices, as in the WT closed form enzyme (Table 3). It could be that the regulatory segment in IDH2 does not need ICT or AKG to fold into an α-helix and successfully close the enzyme. This observation is also aligned with other structural features of IDH2 mutants (differences between cleft widths of the same kind and RMSD values between chains) which are similar to those of the closed form enzymes (Table 7).

**Table 8.** Binding energies of substrates [Kcal/mol] in IDH1 and IDH2 WT and mutant enzymes. The first row indicates the binding site (OC = open cleft; SC = semi-open cleft). The second row contains column headers for binding energy of ICT, AKG, the difference between ICT and AKG (ΔΔ), and the difference between the ΔΔG and the LΔΔ of the binding site (ΔΔΔ). For ΔΔΔ: green: >0; red: <0; white: ≈0. Darker colors highlight values of larger magnitude.

| Structure | OC |  |  |  | SC |  |  |  |
| --- | --- | --- | --- | --- | --- | --- | --- | --- |
| | ICT | AKG | $\Delta\Delta$ | $\Delta\Delta\Delta$ | ICT | AKG | $\Delta\Delta$ | $\Delta\Delta\Delta$ |
| IDH1 WT OF | -8.0 | -5.9 | 2.1 | - | -4.4 | -3.4 | 1.0 | - |
| IDH1 R100Q WS | -4.2 | -2.9 | 1.3 | -0.5 | -4.2 | -3.4 | 0.8 | -0.2 |
| IDH1 R132H WS | -5.3 | -5.0 | 0.3 | -1.5 | -4.7 | -3.0 | 1.7 | 0.7 |
| IDH2 WT OF | -10.7 | -8.9 | 1.8 | - | -6.7 | -3.9 | 2.8 | - |
| IDH2 R140Q WS | -6.7 | -5.6 | 1.1 | -0.7 | -3.9 | -6.3 | -2.4 | -3.4 |
| IDH2 R172K WS | -7.5 | -7.3 | 0.2 | -1.6 | -8.3 | -7.4 | 0.9 | -0.1 |

### Evidence that IDH1 R132H WS and IDH2 R172K WS binding sites are functional

IDH1 R132H WS binding site is similar to IDH1 R132H IF binding site, while IDH2 R172K WS binding site is similar to IDH1 WT CF binding site. These similarities suggest that, upon substrate binding, mutant enzymes behave as WT enzyme with bound substrate in intermediate form and closed form, respectively. The intermediate and closed form correspond to different stages of the enzyme while going from an inactive form into its active form. Therefore, substrate binding on mutant binding sites could be driving the enzymes into their active form.

### The importance of the regulatory segment

Mutant binding sites are deeply influenced by the regulatory segment behavior, which either exposes a previously inaccessibly site (IDH1 R100Q WS and IDH1 R132H WS), is part of the binding site or helps the change in enzyme form (IDH2 R140Q WS and IDH2 R172K WS). However, a regulatory segment like the one in IDH1 has not been previously characterized in enzymes that were not IDH1 homologs. The regulatory segment is in the β-sandwich domain of the enzyme, while the active site is in the Rossman domain (Xu et al., 2004). This means that the regulatory segment is not necessarily exclusive of enzymes with the Rossman domain or enzymes with isocitrate dehydrogenase activity.

## Conclusion

Our methodology, i.e. characterization of the inhibition loss through molecular docking, proved to be a successful approach to our system. Binding energies reported by molecular docking were consistent with the known inhibition/loss of inhibition phenotypes from the different enzymes studied. Additionally, the Jaccard similarity between binding sites allowed structural similarities to be assessed across a set of structures using a defined function, in contrast to traditional methods that require a case by case interpretation. Furthermore, this methodology can be used to explore mutant binding sites from other enzyme families.

Binding sites characterization is one of the most important tasks in protein engineering, since one of the most common goals in protein design is substituting the substrate of an enzyme for another substrate that undergoes a certain chemical reaction of interest.

One of the most recurring questions that researchers seek to answer when studying the IDH1 and IDH2 mutants related to cancer is: Why do the positions IDH1 R132, IDH2 R140 and IDH2 R172 are related to cancer, but IDH1 R100 is not? In this work we propose an explanation, based on a structural characterization, of why the binding site of IDH1 R100Q does not lose its inhibition and therefore it is not related to cancer.

In recent years, several drugs (including some drugs already in clinical research phases) were developed aiming to block cancer-related mutants of IDH1 and IDH2 (Deng et al., 2015; Wu et al., 2015; Zheng et al., 2013). One of the most remarkable aspects of the design of these drugs is the possibility, currently unfulfilled, of targeting several mutations in both enzymes based on the fact that these mutations occur in analogous positions. Our results show that this promiscuous activity is unlikely to be achieved, given that the binding site and the protein structure of different mutant enzymes present significant differences.

